# Sexual interactions and local selection shape reproductive gene diversity and divergence

**DOI:** 10.64898/2026.11.04.763188

**Authors:** Katelyn J. Dykhuizen, Dean M. Castillo

## Abstract

Genes involved in genotype x genotype coevolution display unique patterns of nucleotide diversity and population divergence. In *Drosophila melanogaster, Sex Peptide* is a key sperm competition gene that is an outlier for both π within populations and *Fst* between populations. This pattern is not predicted by current theoretical expectations of selective sweeps highlighting the need to develop predictions to identify genes experiencing balancing selection. We compared π and *F*_*ST*_ in *D. melanogaster* populations to identify other genes following a co-evolutionary model that generates divergence and diversity. Multiple male and female genes that interact with *Sex Peptide* exhibited elevated *F*_*ST*_ *x*π values. Genome-wide analyses also revealed an enrichment of genes critical for neuronal development, pointing to courtship evolution. Specifically, *fruitless*, its interacting genes, and a female behavior gene were identified as outliers. These parallel findings indicate that both sperm competition and mating behaviors are subject to co-evolutionary selection.

## INTRODUCTION

Identifying genes under selection requires specific predictions for what patterns of selection will look like across the genome (Messer & Petrov, 2013; Pritchard et al., 2010). Directional selection produces hard or soft sweeps where an advantageous allele increases inf frequency (Chen et al., 2024; Garud et al., 2021). The two types of sweeps differ in the rate of allele increase and variation surrounding the allele under selection. (Messer & Petrov, 2013; Pritchard et al., 2010). Balancing selection produces a different pattern because alleles are maintained at intermediate frequencies (Ruzicka et al., 2026). Several processes are often classified as balancing selection including heterozygote advantage, environmental tradeoffs, sexual antagonism, temporal or spatially variable selection, and negative frequency dependent selection. The diversity of selection classified as balancing selection makes predicting the exact pattern produced difficult as the specific pattern will likely be different and depend on the timescale and spatial scale that selection operates on (Flintham et al., 2025).

Balancing selection represents a scenario where additional predictions of genomic patterns need to be developed (Johri et al., 2022; Vy et al., 2017). While predictions exist for sexual antagonism/sexual conflict (Amandine et al., 2022; Tosto et al., 2023; Wright et al., 2019; Živkovič et al., 2019) and specific genes involved in negative-frequency dependent selection have been compared to genome wide averages (Chapman et al., 2019), less work has been done to develop and test additional predictions for balancing selection. On reason is that standard tests detect only a subset of loci that have large and symmetrical fitness effects on different fitness components (Brandt et al., 2026). This scenario will not be broadly applicable to many forms of balancing selection even though it can detect some loci under selection (Croze et al., 2017).

The fitness of alleles interacting in genotype x genotype interactions is complex and require a new framework to identify genes under selection. One of the most studied group of genes involved in genotype x genotype interactions are sperm competition genes (Wigby & Chapman, 2004). Sperm competition genes have historically not fit into models of selective sweeps or long-term balancing selection (Kern et al., 2004; Majane et al., 2024; Patlar et al., 2021). This was unexpected, given they are under intense selection, until it was demonstrated that they maintain several alleles through non-transitivity (Clark, 2002; Zhang et al., 2012) and male x female interactions (Clark et al., 1999; Daupagne et al., 2025). Given these mechanisms we would predict genes involved in genotype x genotype interactions would have high levels of nucleotide diversity (π). Since strong co-evolution operates on small spatial scales (Gandon & Michalakis, 2002; Week & Bradburd, 2024) we would also expect few alleles shared between populations resulting in high divergence (*F*_*ST*_). We have observed high π and high *F*_*ST*_ for the sperm competition gene *Sex Peptide* (SP) in *Drosophila melanogaster* (Figure 1) which is known to strongly influence male fitness and co-evolve with female reproductive proteins (Gioti et al., 2012; Hopkins & Perry, 2022; Peng et al., 2024).

**Figure 1.**
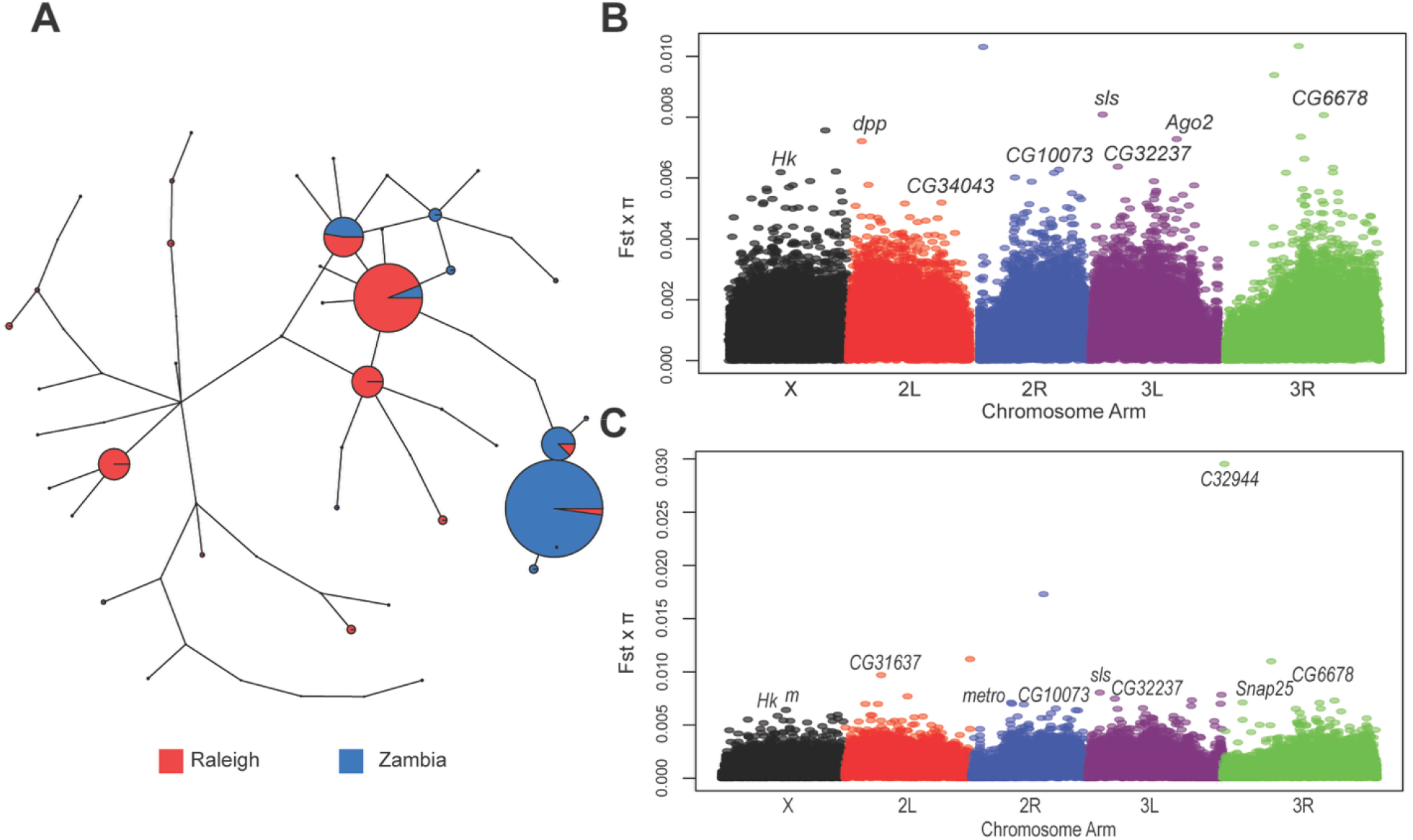
A) The haplotype network for the gene *Sex Peptide (SP)* for Raleigh and Zambia, demonstrating several high frequency haplotypes that differentiate the population, with some shared haplotypes. B). The chromosomal distribution of FST x π outliers for the Raleigh vs Zambia comparison. C). The chromosomal distribution of FST x π outliers for the France vs Zambia comparison. The different colors represent individual chromosome arms that were analyzed with genes labeled that contained outlier SNPs for each chromosome.

This pattern will be unique to genes that evolve through genotype x genotype interactions because π and *F*_*ST*_ are negatively correlated, especially in models of selective sweep (Gillespie, 2000, 2004).We predict that genes co-evolving with genotype x genotype interactions, including sperm competition genes and genes with other roles in sexual interactions, will have both elevated π and *F*_*ST*_. To test this hypothesis we compared genomics patterns of π and *F*_*ST*_ using a composite statistics across populations of *D. melanogaster*.

## METHODS AND MATERIALS

### Genomic Data Sources

Genomic π and *F*_*ST*_ data were accessed from the PopFly database (Hervas et al., 2017) in 1000 base pair windows. 1000 base pair windows were chosen because it is known there are considerable differences in recombination at fine scales within *Drosophila* compared to other systems (Comeron et al., 2012; Fiston-Lavier et al., 2010) and this fine scale results in smaller window of linked selection (Chapman et al., 2019).

### Populations

The first comparison utilized the Raleigh and Zambia populations because of their large sample sizes (n=205 Raleigh, n=197 Zambia). In our initial analysis of Sex Peptide these populations showed a signal of increased π and *F*_*ST*_ for this gene (Fig 1), suggesting we had the power to detect this pattern at additional loci. In the second comparison Zambia and France were used to test the role of geographic distance, by keeping one population constant and comparing to another non-African population that is much closer geographically.

### Identifying Outliers

Our goal was to identify loci that were outliers for both π and *F*_*ST*_. However, we have a priori expectations that there would be a negative correlation between π and *F*_*ST*_ (Gillespie, 2000, 2004), and we observed this for our data (Supplemental Fig 1). In cases of negative dependency composite scores/statistics can be used to capture compensatory extremes (for example an outlier that is 99^th^ percentile in one distribution and 89^th^ in another). In addition, it reduces dimensionality of the data, allowing outliers to stand out more clearly and are used often in genomics (Lotterhos et al., 2017) and other applications such as protein interaction modeling (Eramian et al., 2006). We created a composite score by generating a product distribution and multiplying π and *F*_*ST*_. This allowed us to identify outliers and look for patterns in our subsequent GO analysis. Importantly, previous work has determined that scans using Tajima’s D will only identify a subset of loci under balancing selection (Brandt et al., 2026; Flintham et al., 2025). Since our model makes specific predictions about π and *F*_*ST*_ we needed to use this prediction in our analysis.

### Analyses of a priori sperm competition genes

We determined whether known sperm competition genes showed a pattern of elevated π x *F*_*ST*_ similar to Sex Peptide.. We calculated both an average and a maximum π x *F*_*ST*_ for each gene from (Findlay et al., 2014) since some genes spanned multiple windows. To determine if these values were different than expected by chance, we compared them to a random sample of genomic windows (Chapman et al., 2019). From the genome we subsampled the same number of windows that we did for all sperm competition genes, and then calculated the mean and max for this sample. We repeated this 1000 times to compare this distribution to our observed values for sperm competition genes.

### GO Ontology Analysis

We examined the top 99th percentile windows for π x *F*_*ST*_ to determine which genes were contained within these windows. We then looked for gene ontology (GO) term enrichment of these genes, focusing on terms for biological processes. We used PANGEA (Pathway, Network and Gene-set Enrichment Analysis (Hu et al., 2023). This allowed us to identify other classes of potentially co-evolving genes while testing for biological processes that were overrepresented.

## RESULTS AND DISCUSSION

Looking at genomic wide patterns the X chromosome was underrepresented for outlier loci in both comparisons with 152 outliers for the Raleigh vs Zambia comparison and 87 windows identified as outliers in the France and Zambia comparison (Table 1; Fig 3.). The X chromosome of *D. melanogaster* is often subject to sweeps and reduced variation (Charlesworth et al., 2018), which could be consistent with a lack of balancing selection we have observed. Chromosome 3L was over-represented with 286 outlier windows in Raleigh vs Zambia comparison and 320 France vs Zambia (Table 1). Several inversions differentiate populations of *D. melanogaster*, with inversions on 3R most commonly associated with African and non-African differences (Corbett-Detig & Hartl, 2012; Kapun & Flatt, 2019). These 3R inversions were not overrepresented on our analysis. Interestingly most autosomes aside from 3L show evidence of soft sweeps (Garud et al., 2021), again suggesting that our method is identifying loci that are not consistent with selective sweeps.

**Table 1.** The chromosomal distribution of FST x π outliers for Raleigh (R) and Zambia (Z) comparisons as well as comparisons with France (F). The expected proportion is based on the number of windows per chromosome compared to the total number analyzed.

| Chromosome | Prop. of R vs Z outliers (expected) | Prop. of F vs Z outliers (expected) | Overlap |
|---|---|---|---|
| X | 14.02% (18.84%), n=152 | 7.76% (18.84%), n=87 | 40 |
| 2L | 20.01% (19.33%), n=217 | 24.26% (19.33%), n=272 | 105 |
| 2R | 17.99% (20.26%), n=195 | 19.98% (20.26%), n=224 | 106 |
| 3L | 26.38% (20.62%), n=286 | 28.55% (20.62%), n=230 | 121 |
| 3R | 21.59% (23.44%), n=234 | 19.45% (23.44%), n=218 | 97 |

To understand if these patterns are enriched for specific gene classes we looked at the π x *F*_*ST*_ values for known sperm competition genes and performed a GO analysis on the most extreme outliers.

### Section on a priori sperm candidate genes

We were interested in determining whether sperm competition genes had a similar pattern to the gene Sex Peptide that could suggest co-evolution through genotype x genotype interactions. When looking at both comparisons there were several sperm competition genes had higher maximum π x *F*_*ST*_ values compared to a random sample (Supplemental Table 1; Fig 2).

**Figure 2.**
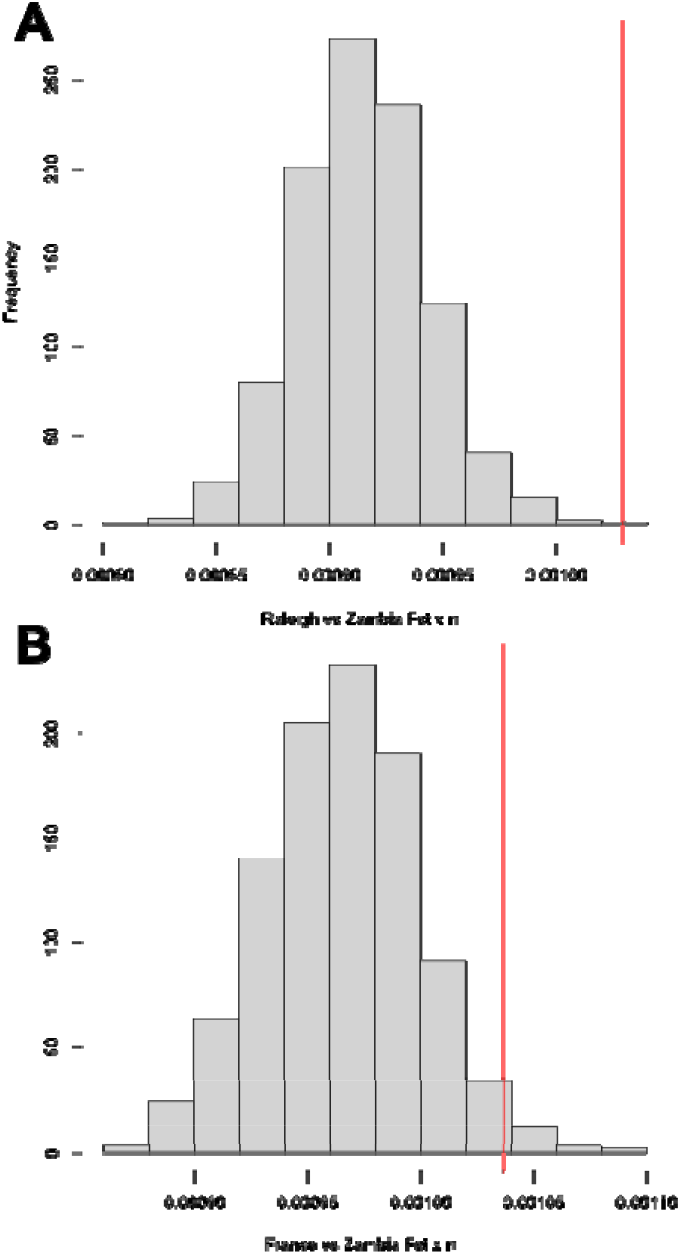
The distribution of mean Fst x π values for a random sample of genomic windows compared with the average value for known sperm competition genes. A) The Raleigh vs Zambia comparison. B) The France vs Zambia comparison. The vertical line represents the average max window value across all sperm competition genes.

This suggests that there is a unique aspect of the biology of sperm competition genes that produces this pattern. Several male sperm competition genes were outliers including *Acp53Ea* and *CG9997* as well as female specific genes *Sex Peptide Receptor (SPR)* and *hadley*. In contrast, several sperm competition genes show little variation with low π and low *F*_*ST*_. For a complex trait such as sperm competition, where there are hundreds of reproductive proteins produced by both males and females, the number of genes that are targets of selection can be difficult to determine (Garlovsky et al., 2024; Humphries et al., 2008; Lymbery et al., 2018). It is possible that only a few of the deeply connected proteins respond to genotype x genotype co-evolution, whereas the others do not. Consistent with this, *Sex Peptide* and *Sex Peptide Receptor* have a close co-evolutionary history (Hopkins & Perry, 2022; Peng et al., 2024) and had the largest π x *F*_*ST*_ for any of the male or female genes.

### Identifying co-evolving reproductive genes

To determine if reproductive processes are overresponded in our outliers we conducted a GO analysis on the most extreme outliers for both comparisons. The most significantly enriched biological term for both Raleigh and Zambia comparisons and France and Zambia comparisons was neuron differentiation (Supplemental Table 2). Other top terms were also related to neuron development. Sperm competition was not a significant GO term, but this is not surprising given the small number of genes that are associated with this term (n=25 compared to n=864 for neuron differentiation).

The identification of neuronal genes could suggest co-evolution through genotype interactions involved in courtship behavior. For example, *fruitless (fru)* contributed to many of the significant GO terms. While *fru* is essential for neuronal development it is most often discussed in terms of sexual behaviors, specifically male courtship (von Philipsborn et al., 2014; Yamamoto & Koganezawa, 2013). Previous work has suggested *fru* experiences balancing selection caused by antagonistic pleiotropy (Jardine et al., 2021). This signal was identified using polymorphism data and Tajima’s D and it was found that two alleles are maintained in both

African and North American populations. Antagonistic pleiotropy would not be responsible for the high *F*_*ST*_ that we have observed. While increased π can be caused by pleiotropy this did not drive the signal alone as *F*_*ST*_ for *fru* was above the 90th percentile for genomic *F*_*ST*_.

Similar to the case of the sperm competition genes, multiple genes with known behavioral effects were outliers and importantly some of these genes interact with *fru*. The gene *stripe (sr)* has a GO term of neuronal differentiation and has been connected to male courtship behavior. *stripe* has been shown to be transcriptionally active in *fru* expression neurons when males sexually interact with females (Takayanagi-Kiya et al., 2023). Similarly, *big brain (bib)* interacts with *fru* and can enhance expression of *fru* in sexually dimorphic brain regions (Goto et al., 2011). The shared pattern for these interacting genes demonstrates the ability of this framework to identify networks related to reproductive behaviors. We also observed female behavior genes contributing to GO enrichment. The gene *alan shepard* has known roles in female mating behavior and female mate preference (Chen et al., 2014; Roy & Castillo, 2024).

While there is no indication of co-evolution between *shep* and *fru* directly, male traits and female mate preferences have correlated patterns of evolution between populations of *D. melanogaster* (Ferveur, 2010; Grillet et al., 2012; Jin et al., 2022).

Taken together these analyses have identified a population genetic pattern that is consistent with a genotype x genotype co-evolutionary model. The *a priori* sperm competition genes interact in a network and displayed high nucleotide diversity within populations and high differentiation between populations. Behavioral genes that are known to interact showed a similar pattern suggesting they experience genotype x genotype co-evolution. This expectation can be examined in other cases where co-evolution might be expected, and could also be used to identify genes that may underlie co-evolving phenotypes as we have demonstrated.

