## Supplemental Materials for "Sexual interactions and local selection shape reproductive gene diversity and divergence"

**Supplemental Material**

**Supplemental Table 1.** The summary of Fst x π statistics for genes with known roles in sperm competition. The location of the gene and the number of windows the gene spans are included with the average percentile and max percentile for both comparisons.

| Chromosome | Gene | Number of windows | R vs Z Average Percentile | R vs Z Max Percentile | F vsZ Average Percentile | F vsZ Max Percentile |
| --- | --- | --- | --- | --- | --- | --- |
| 2L | Accessory gland protein 29AB | 2 | 92.43 | 95.42 | 95.45 | 98.14 |
| 2L | Polycystic kidney disease 2 | 4 | 74.82 | 85.63 | 79.96 | 95.03 |
| 2L | Accessory gland protein 26Aa | 3 | 83.40 | 92.31 | 89.95 | 94.32 |
| 2L | Accessory gland protein 36DE | 4 | 33.32 | 83.51 | 34.81 | 84.37 |
| 2R | Accessory gland protein 53Ea | 7 | 98.53 | 99.94 | 98.64 | 99.95 |
| 2R | lectin-46Cb | 3 | 70.24 | 87.03 | 65.05 | 85.56 |
| 2R | antares | 1 | 85.23 | 85.23 | 89.17 | 89.17 |
| 2R | CG17575 | 3 | 80.80 | 97.87 | 72.68 | 96.42 |
| 2R | CG1652 | 3 | 70.24 | 87.03 | 65.05 | 85.56 |
| 3L | Sex peptide | 1 | 99.85 | 99.85 | 99.63 | 99.63 |
| 3L | dynein, axonemal, heavy chain 3 | 48 | 70.75 | 99.98 | 60.85 | 98.94 |
| 3L | Esterase 6 | 1 | 43.98 | 43.98 | 41.90 | 41.90 |
| 3L | Accessory gland protein 62F | 9 | 78.39 | 94.68 | 78.40 | 92.77 |
| 3L | dynein, axonemal, intermediate chain 2 | 2 | 38.18 | 49.60 | 51.20 | 56.99 |
| 3L | Seminase | 50 | 15.23 | 66.94 | 15.96 | 59.61 |
| 3R | Abdominal B | 62 | 60.91 | 99.03 | 54.57 | 97.58 |
| 3R | MSA micropeptide | 90 | 65.21 | 98.82 | 64.83 | 98.51 |
| 3R | CG9997 | 1 | 94.91 | 94.91 | 94.08 | 94.08 |
| 3R | Neprilysin 2 | 16 | 9.48 | 32.96 | 9.18 | 28.96 |
| 3R | intrepid | 3 | 72.13 | 90.37 | 80.42 | 91.15 |
| 3R | Glucose dehydrogenase | 18 | 23.74 | 62.48 | 24.80 | 65.57 |
| 3R | aquarius | 1 | 30.64 | 30.64 | 22.15 | 22.15 |
| 3R | lost boys | 33 | 76.45 | 99.37 | 79.12 | 98.92 |
| 3R | hadley | 20 | 80.86 | 99.21 | 76.45 | 98.95 |
| 3R | Esp | 11 | 82.81 | 96.67 | 75.52 | 91.41 |
| X | fra mauro | 3 | 81.35 | 96.48 | 82.24 | 95.45 |
| X | SPR | 65 | 80.64 | 99.76 | 80.87 | 99.41 |

**Supplemental Table 2.** The top three GO terms from the Raleigh vs Zambia Fst x π outliers as an example of significant enrichment for genes important in neuron differentiation and development. We have provided the list of the genes for these specific GO categories.

| GO term | Number of genes | FDR | Names of Genes |
| --- | --- | --- | --- |
| neuron differentiation | 79 | 0.0004118 | mts,Ptp99A,Fur1,gl,fru,cdi,robo1,Con,hdc,sqz,Spn,Sema2a,fra,Mad,wts,klu,Rab11,ara,Pvf3,nub,sif,Snoo,chinmo,futsch,bun,Mob2,Ca-beta,Mp,NT1,wb,tinc,cnc,hts,dally,Sema2b,ab,PIG-S,bib,dpp,EcR,E(z),for,fz,klar,mam,pnt,ru,shn,tkv,Ten-a,ttv,mbl,vri,fry,Pka-R2,melt,caps,stan,gukh,beat-Ic,beat-Ib,daw,beat-IIIc,Smyd4-3,Dg,Efhc1.2,Cip4,RhoGEF64C,Pura,tey,Archease,Sar1,DIP-gamma,tai,robo3,exex,Trim9,CG32373,shep |
| neuron development | 65 | 0.0014417 | fry,Pka-R2,caps,stan,gukh,beat-Ic,beat-Ib,daw,beat-IIIc,Smyd4-3,Dg,EcR,E(z),for,fz,klar,pnt,shn,Pvf3,nub,sif,Snoo,chinmo,futsch,Mob2,Ca-beta,Mp,NT1,wb,tinc,cnc,hts,dally,Sema2b,ab,PIG-S,ttv,Ten-a,mbl,Ptp99A,Fur1,fru,robo1,Con,hdc,sqz,Spn,Sema2a,fra,wts,Rab11,ara,Efhc1.2,RhoGEF64C,Pura,tey,Archease,Sar1,DIP-gamma,tai,robo3,exex,Trim9,CG32373,shep |
| generation of neurons | 82 | 0.0018887 | Efhc1.2,Cip4,RhoGEF64C,Pura,tey,Archease,Sar1,DIP-gamma,CG7834,tai,robo3,exex,Trim9,CG32373,shep,bib,dpp,EcR,E(z),for,fz,klar,mam,pnt,ru,shn,tkv,ttv,Ten-a,mbl,wake,vri,fry,Pka-R2,melt,caps,stan,gukh,beat-Ic,beat-Ib,daw,beat-IIIc,Smyd4-3,Dg,clu,mts,Ptp99A,Fur1,gl,fru,cdi,robo1,Con,hdc,sqz,Spn,Sema2a,fra,Mad,wts,klu,Rab11,ara,Pvf3,nub,sif,Snoo,chinmo,futsch,bun,Mob2,Ca-beta,Mp,NT1,wb,tinc,cnc,hts,dally,Sema2b,ab,PIG-S |

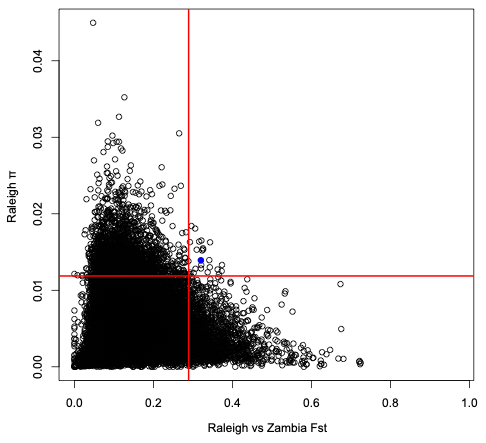

**Supplemental Figure 1.** The distributions of Fst and π for the Raleigh vs Zambia comparison demonstration the lack of genes that are outliers for both variables. The red lines highlight the 90th percentiles.

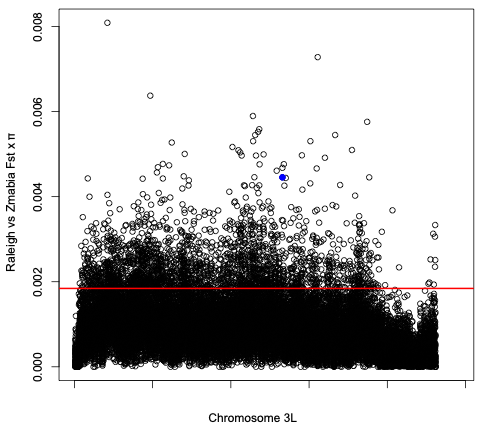

**Supplemental Figure 2.** The distributions of Fst and π for the Raleigh vs Zambia comparison demonstration the lack of genes that are outliers for both variables. The red lines highlight the 90th percentiles.
